# Transcriptome Analysis of Four *Arabidopsis thaliana* Mediator Tail Mutants Reveals Overlapping and Unique Functions in Gene Regulation

**DOI:** 10.1101/309153

**Authors:** Whitney L. Dolan, Clint Chapple

**Affiliations:** Department of Biochemistry, Purdue University, West Lafayette, IN 47907, USA and Purdue Center for Plant Biology, West Lafayette, IN 47907, USA.

**Author notes:** **Reference numbers**: Gene expression data are available in the Gene Expression Omnibus under accession GSE95574. **Corresponding author**: Clint Chapple, Department of Biochemistry, Purdue University 175 South University St., West Lafayette, IN 47907.

**Keywords:** Mediator, Arabidopsis, transcription regulation, gene expression

## Abstract

The Mediator complex is a central component of transcriptional regulation in Eukaryotes. The complex is structurally divided into four modules known as the head, middle, tail and kinase modules, and in *Arabidopsis thaliana*, comprises 28-34 subunits. Here, we explore the functions of four Arabidopsis Mediator tail subunits, MED2, MED5a/b, MED16, and MED23, by comparing the impact of mutations in each on the Arabidopsis transcriptome. We find that these subunits affect both unique and overlapping sets of genes, providing insight into the functional and structural relationships between them. The mutants primarily exhibit changes in the expression of genes related to biotic and abiotic stress. We find evidence for a tissue specific role for MED23, as well as in the production of alternative transcripts. Together, our data help disentangle the individual contributions of these MED subunits to global gene expression and suggest new avenues for future research into their functions.

## INTRODUCTION

The Mediator complex is an essential co-regulator of eukaryotic transcription, participating in many of the events surrounding transcription initiation (Poss *et al*. 2013; Allen and Taatjes 2015). Mediator bridges the divide between enhancer-bound transcription factors and promoter-bound RNA Polymerase II (Pol II) to facilitate assembly and function of the preinitiation complex. The individual subunits of the complex have been assigned to four modules, known as the head, middle, tail, and kinase modules, based on their positions within the complex (Figure 1). The head and middle modules contact Pol II, while the tail module primarily interacts with transcription activators (Figure 1; Tsai et al., 2017; Jeronimo et al., 2016; Lee et al., 1999; Myers et al., 1999; Park et al., 2000; Zhang et al., 2004). The kinase module reversibly associates with the rest of the complex and is thought to play a negative regulatory role by inhibiting interaction of Mediator with Pol II (Elmlund *et al*. 2006; Knuesel *et al*. 2009; Tsai *et al*. 2013). The core Mediator complex has recently been redefined as just the middle and head modules as they are the minimal components required for Mediator to stimulate transcription (Cevher *et al*. 2014; Plaschka *et al*. 2015; Jeronimo *et al*. 2016). Although the core is capable of functioning independently, the majority of evidence suggests that the tail is associated with the core under most circumstances.

**Figure 1.**
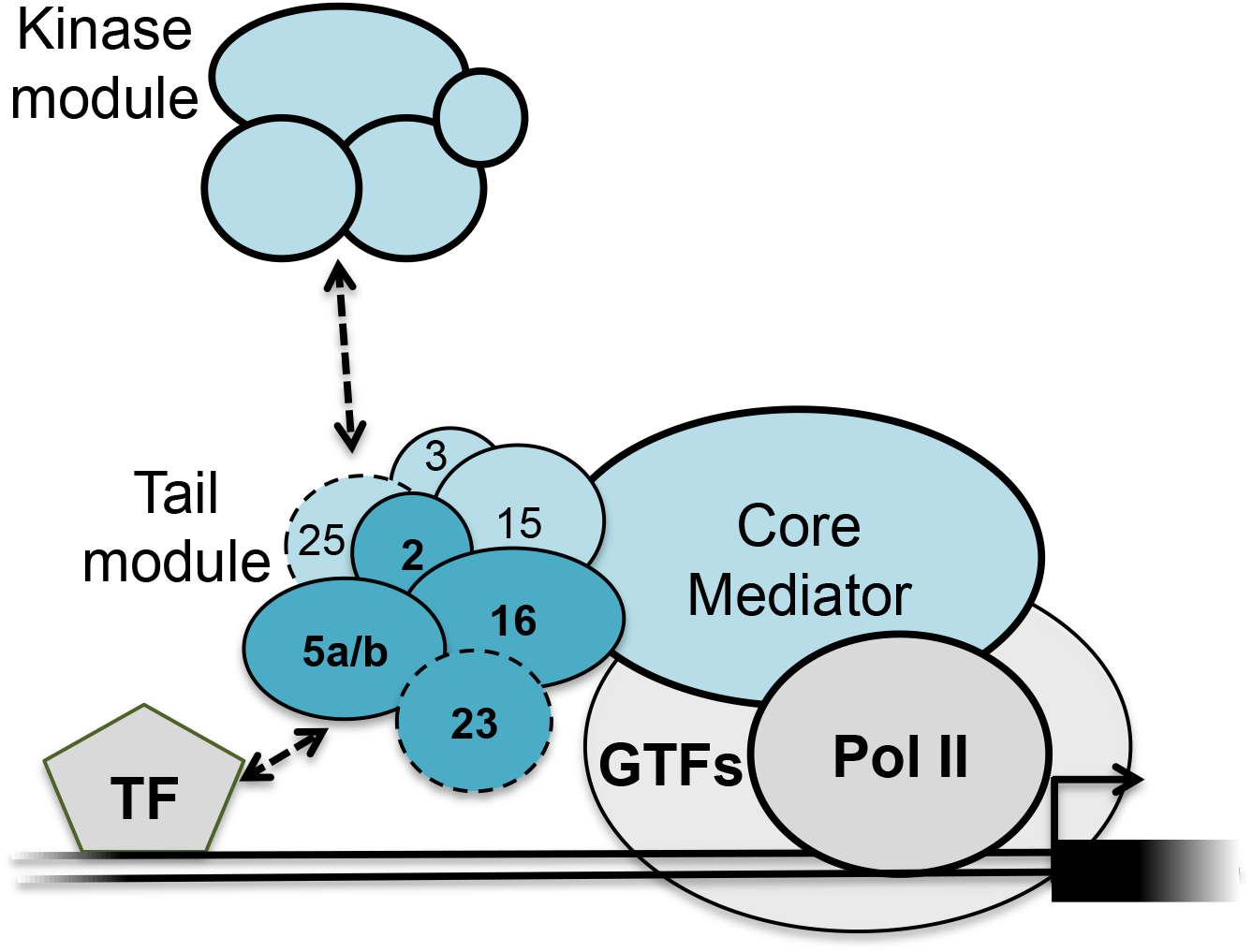
Model of the Arabidopsis Mediator complex. Core Mediator interacts with RNA Pol II and the general transcription factors (GTFs). The tail module (numbered subunits) interacts with DNA-bound transcription factors (TF and the dissociable kinase module. Dark blue subunits are those studied here. The positions of the subunits outlined with dashed lines are not well determined.

Given that the middle and head modules can be recruited to promoters and facilitate preinitiation complex (PIC) assembly independent of the tail module, it appears that a major role of the tail is to increase the probability of Mediator-PIC interactions by recruiting and tethering the complex to promoter-proximal transcription factors (Jeronimo *et al*. 2016); however, this does not appear to be the only role of the tail and many questions remain regarding its structure and function. The tail is highly flexible and has thus been difficult to visualize using the composite cryo-EM imaging techniques that have recently enabled high resolution structures of core Mediator (Tsai *et al*. 2017). In addition, many studies of Mediator structure have focused on yeast Mediator complexes, which lack some tail subunits found in humans and plants (Bourbon 2008). Structural, genetic, and functional data from a number of organisms support the existence of two submodules within the tail, one comprising MED2, MED3, and MED15, and another comprising MED5, MED16, and MED23 (Li *et al*. 1995; Ito *et al*. 2002; Zhang *et al*. 2004; Béve *et al*. 2005; Robinson *et al*. 2015). Although loss of MED16 results in separation of the rest of the tail from the complex, the free MED2-MED3-MED15 submodule can still be recruited by transcription factors to activate transcription (Zhang *et al*. 2004; Galdieri *et al*. 2012). Aside from its role in recruiting Mediator to promoters, the tail module also facilitates reinitiation by helping to maintain a scaffold PIC (Reeves and Hahn 2003). Negative regulation of transcription also occurs through the tail in some instances. CDK8, the enzymatically active subunit of the kinase module, has been shown to phosphorylate both MED2 and MED3, resulting in gene repression (van de Peppel *et al*. 2005; Gonzalez *et al*. 2014).

In Arabidopsis, Mediator tail subunits have been shown to be required for the regulation of a variety of processes (reviewed in Yang et al., 2015; Samanta and Thakur, 2015). Mediator tail subunits MED16 and MED25 are two of the most extensively studied Arabidopsis MED subunits. *MED16/SFR6* was first identified for its role in freezing tolerance and *MED25/PFT1* for its role in flowering (Knight 1999; Cerdán and Chory 2003). Since then, both have been shown to function extensively in the regulation of defense-related genes, as well as a number of other processes (reviewed in Dolan and Chapple 2017). MED2 has been less well studied, but has been shown to share some functions with MED14 and MED16 in cold-regulated gene expression (Hemsley *et al*. 2014). MED5a/b also share some functions with MED14, and MED16, but in this case in the regulation of dark induced gene expression (Hemsley *et al*. 2014). From these studies and others it has become increasingly apparent that normal gene expression requires the concerted action of multiple MED subunits, making it difficult to disentangle the functions of individual subunits. This fact was highlighted by the observation that nine different MED subunits are required for methyl-jasmonate induced expression of *PDF1.2* (Wang *et al*. 2016).

Previously, we showed that MED2, MED16, and MED23 are differentially required for the function of *ref4-3*, a semi-dominant *MED5b* mutant that negatively impacts phenylpropanoid accumulation (Dolan et al. 2017). In the present study, we explore the effects of disrupting *MED2, MED5a/b, MED16*, and *MED23* on genome-wide transcription to gain a broader understanding of their roles in gene regulation and their functional relationships to one another. As expected, we find that these subunits have both distinct and overlapping roles in gene regulation. These data lay a foundation for teasing apart the individual contributions of these MED subunits to the expression of different pathways and genes, and more importantly, for understanding how they function as a unit.

## RESULTS

### The ***med*** tail mutants exhibit minor changes in development

Previously, we isolated T-DNA lines of *MED2, MED5a, MED5b, MED16*, and *MED23*, and showed that full-length transcripts of the genes in which the insertions are located are either abolished or substantially reduced (Dolan *et al*. 2017). Using the *med5a* and *med5b* mutants, we created a *med5ab* double mutant, as the two genes appear to be largely interchangeable within the complex (Bonawitz et al. 2012). Under our growth conditions, all of these *med* mutants develop similarly to wild-type plants (Figure 2 and Dolan et al. 2017), with a few exceptions. The *med2* plants fail to stand erect as they get taller, indicating that they have weakened inflorescences (Dolan *et al*. 2017). In addition, the *med2* and *med16* mutants appear to have slightly smaller rosettes than wild type early on in development (Figure 2). We also observed that *med2* and *med5ab* plants flower early (discussed in more detail below), whereas *med16* is known to be late-flowering (Knight *et al*. 2008).

**Figure 2.**
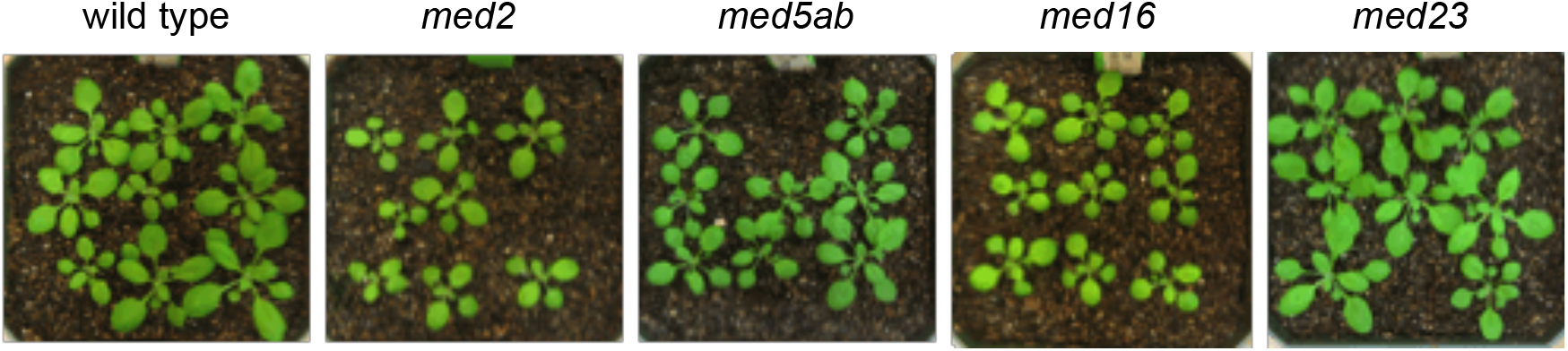
The *med* mutants grow similar to wild-type plants. A representative subset of the 18-day old plants used for RNAseq.

### ***med*** tail mutants have unique effects on the transcriptome

We grew all of the *med* mutants under a 16 hr light, 8 hr dark cycle for 18 days, at which time we collected whole-rosettes for RNA extraction followed by RNAseq analysis (Figure 2 and Dolan et al. 2017). In our previous analysis of the data, we showed that in the *med2* and *med5ab* mutants, significantly more genes are downregulated than upregulated (Dolan et al. 2017). We also showed that many more genes are differentially expressed in *med16* than in the other mutants, with a similar number of genes being up– or downregulated, and that in *med23* very few genes are differentially expressed (Dolan et al. 2017). Here, we analyzed the same data in more detail using a stricter false discovery rate (FDR) of 0.01 and the same 2-fold change minimum cutoff, with the expectation that these genes would be more likely to represent those that directly require the MED subunits for their proper expression (Figure 3). Using these criteria we found that there were 364 differentially expressed genes (DEGs) in *med2* (53**↑**, 311 **↓**), 305 DEGs in *med5ab* (66**↑**, 239**↓**), 768 DEGs in *med16* (289**↑**, 479**↓**), and 47 DEGs in *med23* (15**↑**, 33**↓**).

**Figure 3.**
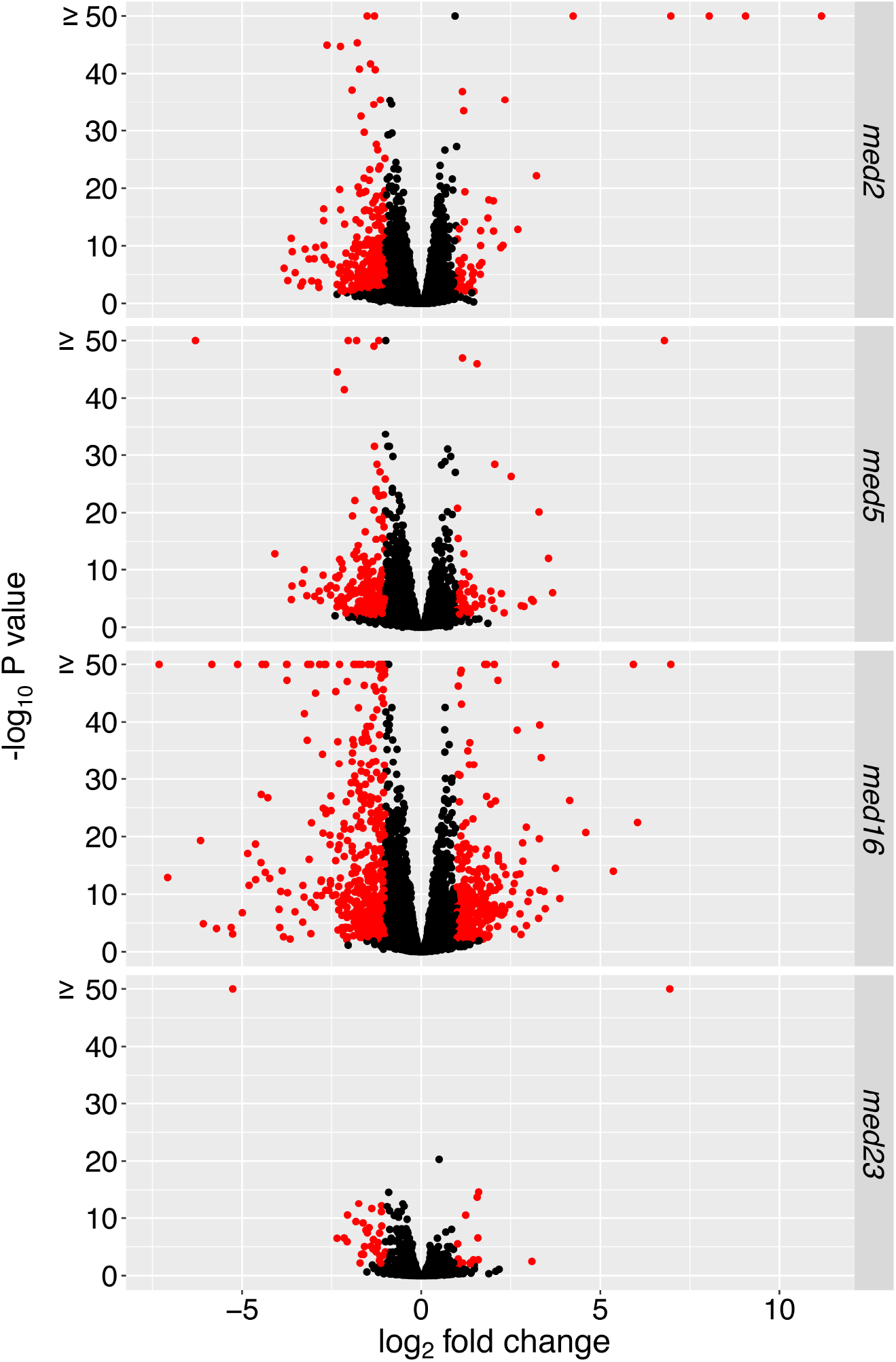
Volcano plots showing differential gene expression in the *med* mutants. Genes with an adjusted *P* value of < 0.01 and a log_2_ fold-change ≥1 are highlighted in red.

Comparison of the genes that were differentially expressed in each of the four *med* mutants showed that there were a large number of DEGs unique to each line, except for *med23*. Most of the DEGs in *med23* were also differentially expressed in *med5ab* and/or *med16* (Figure 4). There were also a large number of genes (103) that were shared only by *med2* and *med16*.

**Figure 4.**
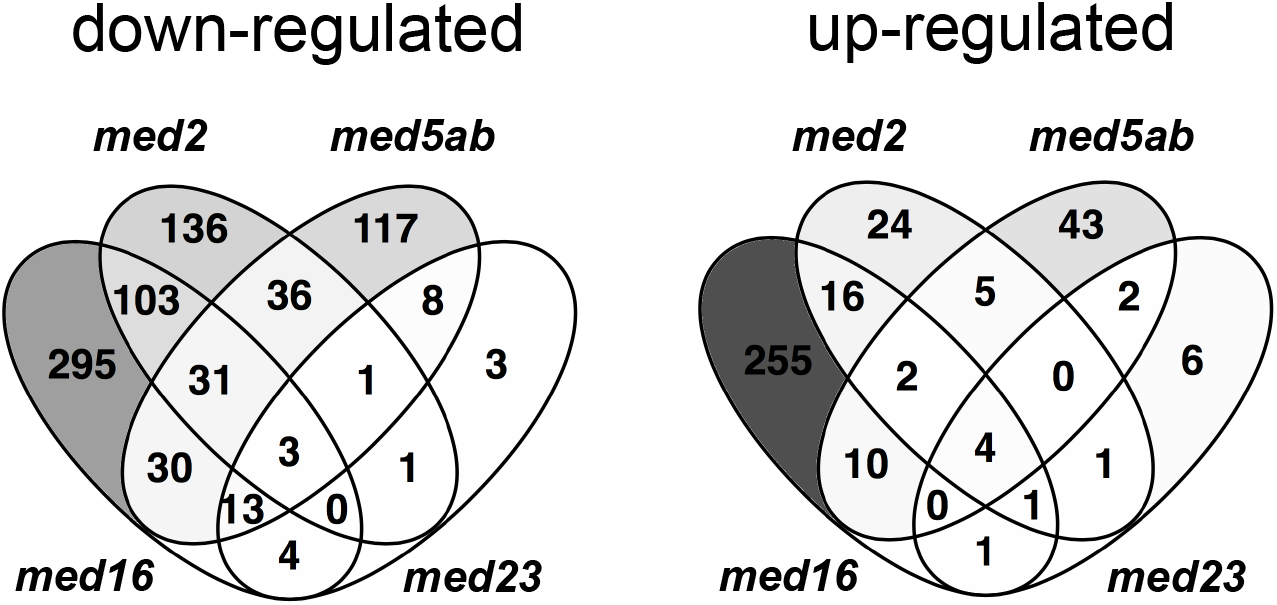
Overlap in up– or downregulated genes between the *med* mutants. Includes all genes that were differentially expressed compared to wild type (FDR <0.01) with an absolute log_2_ fold change > 1.

Only three genes were downregulated in all four mutants. They are *DRM2*, which encodes an auxin/dormancy associated protein, *ERF105*, which encodes an ethylene responsive transcription factor, and *AT1G35210*, which encodes a hypothetical, chloroplast localized protein. Similarly, only four genes were upregulated in all four mutants. They include one gene from the copia-like retrotransposon family (AT5G35935), one gene from the gyspy-like retrotransposon family (AT5G28335), the 5.8S rRNA gene (AT3G41979), and a gene that encodes a defensin-like family protein (AT2G16367). Given that there were so few DEGs in *med23*, it was not surprising that there was so little overlap between all four mutants. For this reason, we also looked at the DEGs shared just by *med2, med5ab*, and *med16* (Table 1). Among the 31 genes that are downregulated in the three mutants, there is no significant enrichment of any gene ontology (GO) terms. The three mutants share only two upregulated genes, those being *MYB47* and *SAUR12*.

**Table 1.** Genes that are differentially expressed in *med2, med5ab* and *med16* mutants.

| AGI | Gene description | log<sub>2</sub> fold-change |  |  |
| --- | --- | --- | --- | --- |
|  |  | med2 | med5ab | med16 |
| up-regulated |  |  |  |  |
| AT1G18710 | myb domain protein 47 (MYB47) | 1.69 | 1.49 | 3.31 |
| AT2G21220 | SAUR-like auxin-responsive protein family (SAUR12) | 1.66 | 1.34 | 1.62 |
| down-regulated |  |  |  |  |
| AT1G10070 | branched-chain amino acid transaminase 2 (BCAT-2) | -1.57 | -1.28 | -1.44 |
| AT1G15125 | S-adenosyl-L-methionine-dependent methyltransferases superfamily protein | -1.52 | -2.19 | -4.47 |
| AT1G19380 | Protein of unknown function (DUF1195) | -1.31 | -1.12 | -1.77 |
| AT1G21100 | O-methyltransferase family protein | -1.14 | -1.21 | -1.63 |
| AT1G27020 | unknown protein | -1.56 | -1.25 | -1.16 |
| AT1G51820 | Leucine-rich repeat protein kinase family protein | -1.32 | -2.05 | -1.44 |
| AT1G69880 | thioredoxin H-type 8 (TH8) | -3.24 | -1.49 | -3.30 |
| AT1G73330 | drought-repressed 4 (DR4) | -1.72 | -1.15 | -2.66 |
| AT2G05440 | GLYCINE RICH PROTEIN 9 (GRP9) | -1.47 | -4.08 | -4.35 |
| AT2G26560 | phospholipase A 2A (PLA2A) | -1.74 | -1.91 | -2.53 |
| AT2G40330 | PYR1-like 6 (PYL6) | -1.59 | -1.05 | -1.80 |
| AT2G43120 | RmlC-like cupins superfamily protein | -1.93 | -1.10 | -3.76 |
| AT3G10020 | unknown protein | -1.09 | -1.14 | -1.51 |
| AT3G22060 | Receptor-like protein kinase-related family protein | -1.59 | -1.26 | -1.23 |
| AT3G26200 | CYP71B22 | -2.25 | -1.17 | -3.74 |
| AT3G43828 | CACTA-like transposase family | -1.72 | -1.25 | -1.42 |
| AT3G48520 | CYP94B3 | -3.83 | -1.92 | -3.97 |
| AT3G49620 | DARK INDUCIBLE 11 (DIN11); 2-oxoglutarate (2OG) and Fe(II)-dependent oxygenase | -2.29 | -1.61 | -1.74 |
| AT3G50010 | Cysteine/Histidine-rich C1 domain family protein | -1.07 | -2.29 | -2.67 |
| AT3G51400 | protein of unknown function (DUF241) | -1.80 | -1.01 | -2.26 |
| AT4G11460 | cysteine-rich RLK (RECEPTOR-like protein kinase) 30 (CRK30) | -1.71 | -1.09 | -1.11 |
| AT4G15210 | beta-amylase 5 (BAM5) | -1.59 | -3.26 | -4.62 |
| AT4G33467 | unknown protein | -1.98 | -2.74 | -4.47 |
| AT4G35770 | SENESCENCE 1; Rhodanese/Cell cycle control phosphatase superfamily protein | -1.54 | -1.76 | -2.15 |
| AT5G14360 | Ubiquitin-like superfamily protein | -1.71 | -1.41 | -1.70 |
| AT5G39890 | Protein of unknown function (DUF1637) | -1.99 | -1.14 | -1.66 |
| AT5G41761 | unknown protein | -1.14 | -1.91 | -4.84 |
| AT5G44420 | plant defensin 1.2 (PDF1.2) | -2.16 | -2.85 | -7.08 |
| AT5G51790 | basic helix-loop-helix (bHLH) DNA-binding superfamily protein | -1.85 | -1.08 | -1.27 |
| AT5G56870 | beta-galactosidase 4 (BGAL4) | -1.06 | -1.04 | -1.05 |
| AT5G62360 | Plant invertase/pectin methylesterase inhibitor superfamily protein | -1.04 | -1.92 | -4.44 |

To determine how the expression profiles of the mutants correlate more broadly, we compared the expression of all DEGs that had an FDR < 0.01 in at least one of the mutants (Figure 5A). This approach revealed a positive correlation in the expression profiles of *med5ab* and *med23* (*r* = 0.61). There was little correlation between the other expression profiles with *med5ab* and *med16* being the most different from one another. Stronger correlations were observed when the comparisons were limited to only those genes that met the FDR cutoff in both mutants (Figure 5B), except in the case of *med16* and *med5ab*, in which many genes were differentially expressed in opposite directions.

**Figure 5.**
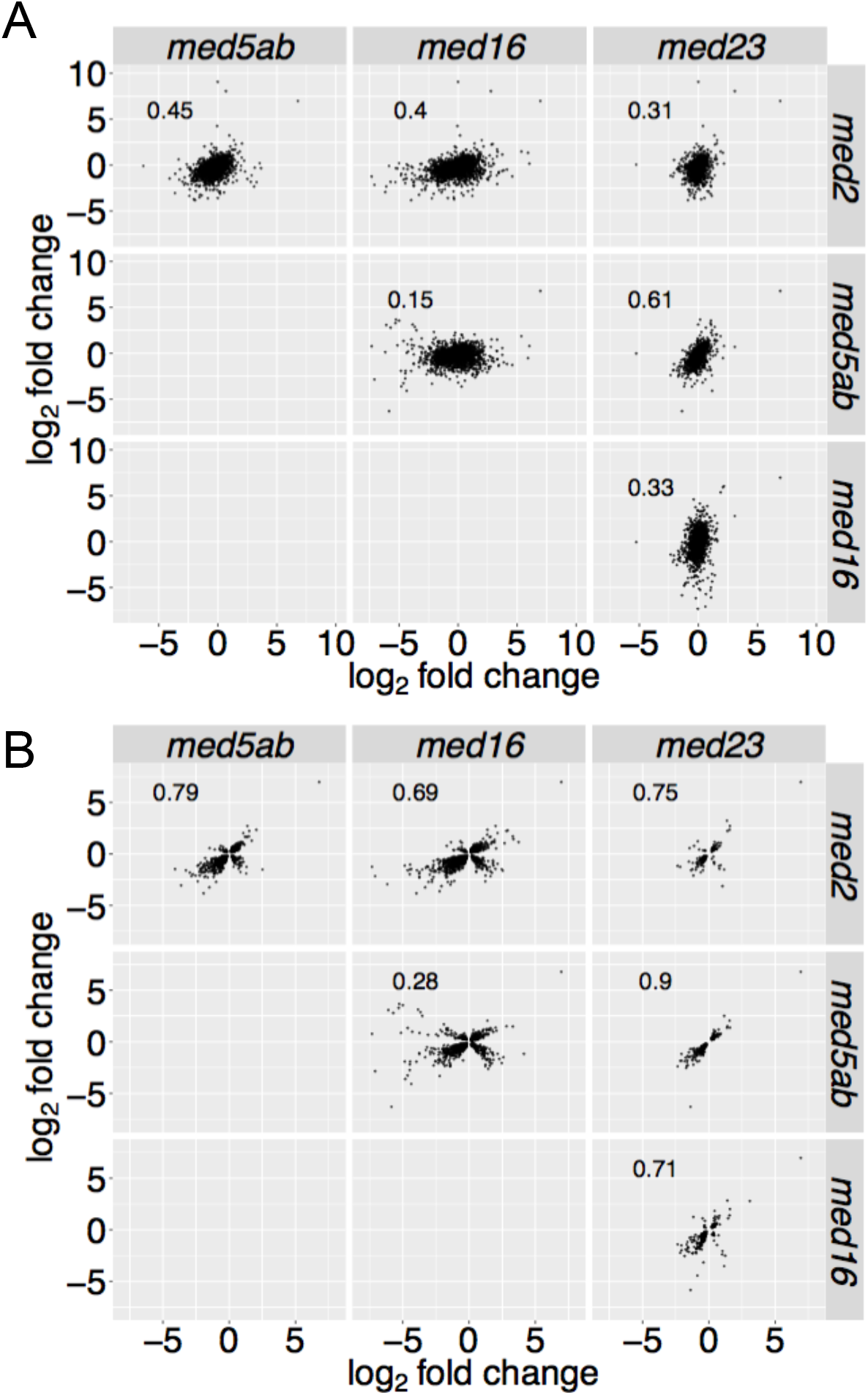
Pairwise comparison of the gene expression profiles of the *med* mutants. Scatter plots comparing the log_2_ fold change in expression compared to wild type of genes that are (A) differentially expressed in any of the four mutants (FDR < 0.01) or differentially expressed in both mutants being compared. The Pearson (r) correlation is given for each pair of comparisons.

### MED tail mutants affect different biological processes

Gene ontology (GO) term enrichment analysis of the DEGs in each of the mutants showed substantial differences in the pathways and processes affected (Figure 6). Defense and cellular stress pathways are upregulated in *med16*, whereas the same pathways are downregulated in *med5ab*. Several other defense pathways are downregulated in *med5ab* that are not affected in the other mutants, as are “vasculature development” and “response to cold”.

**Figure 6.**
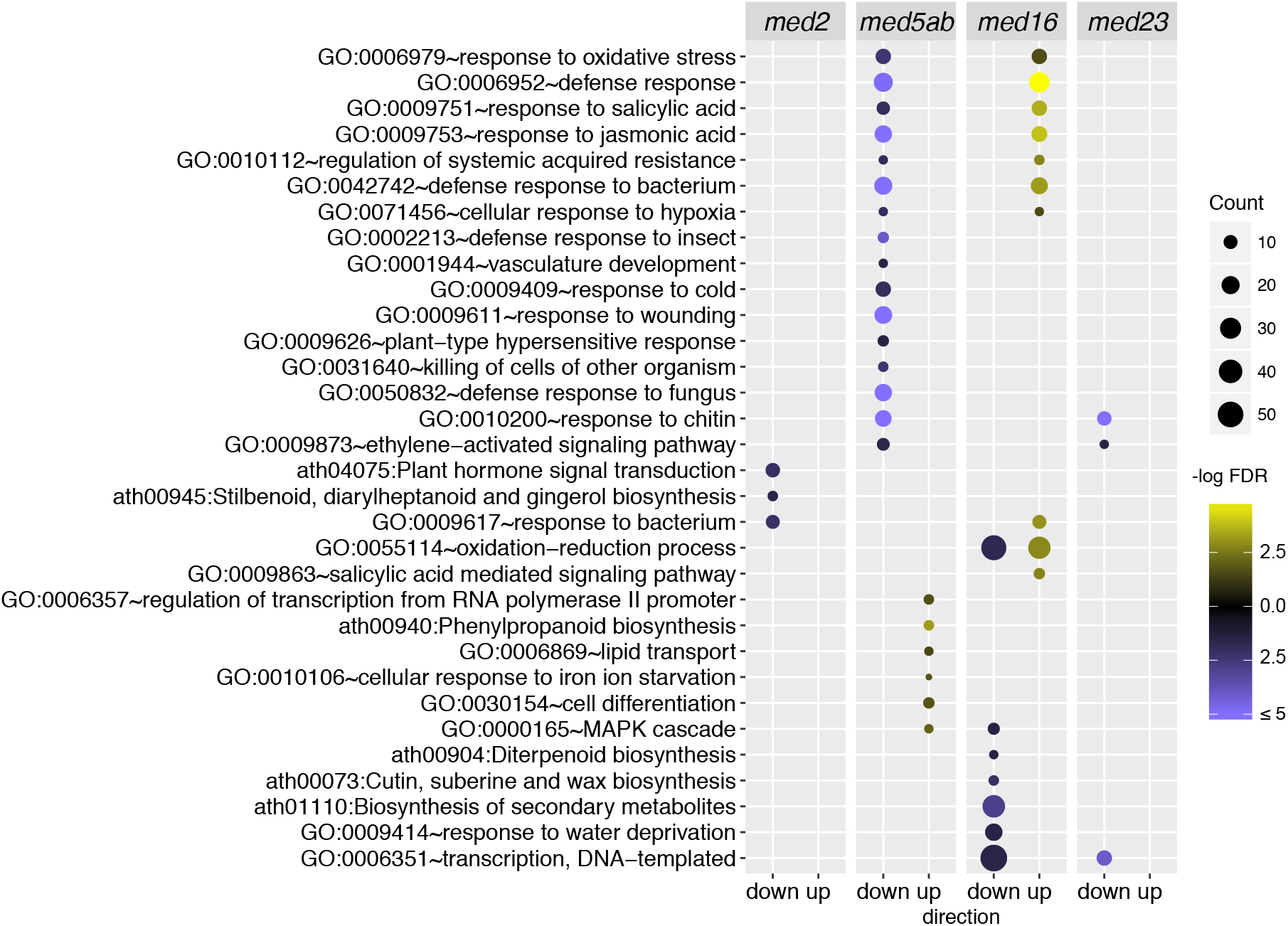
Gene ontology enrichment among genes that are differentially expressed in the *med* mutants. Enrichment of “Biological process” GO-terms and KEGG pathways. Terms that were largely redundant were removed. Direction indicates the subset of genes with increased or decreased expression in each of the mutants compared to wild type (FDR < 0.01, absolute log_2_ fold change > 1). The brightness of the circles indicates the significance of the term or pathway (-log FDR) and their size indicates the number of genes that are associated with that term or pathway.

In *med16*, many genes related to the biosynthesis of secondary metabolites, response to water deprivation, and transcription, are downregulated. Among the 311 genes that are downregulated in *med2*, only three GO-terms are enriched; they are, “plant hormone signal transduction”, “response to bacterium”, and “stillbenoid, diarylheptanoid and gingerol biosynthesis”. Likewise, only three GO-terms are enriched in the *med23* mutant and all are downregulated. They include “response to chitin” and “ethylene-activated signaling pathway”, which are shared with *med5ab*, and “transcription, DNA-templated”, which is shared with *med16*.

### Hierarchical clustering identifies genes that require different subsets of MED tail subunits for their proper expression

To identify groups of genes that behave similarly or differently in the *med* mutants, we performed hierarchical clustering using the complete set of 1080 DEGs (Figure 7). Six major gene clusters were identified (Figure 7A and B). Cluster 1 contains genes that are largely downregulated in *med2* and *med5ab* and is enriched for defense-related genes (Figure 7C, Table 2). Cluster 2 contains genes that are downregulated in all of the mutants and is enriched for genes related to water deprivation and hormone signal transduction (Figure 7C, Table 2. Cluster 3 contains genes that are downregulated in *med16* and to some extent, *med2* (Figure 7C, Table 2). Cluster 3 genes encode proteins involved in secondary metabolite biosynthesis, transcription regulation, and extracellular processes. Cluster 4 contains genes that are upregulated in *med16* and downregulated in *med2* and *med5ab* and is enriched for genes involved in numerous defense pathways (Figure 7C, Table 2). Cluster 5 contains genes that are upregulated in all of the mutants and is enriched for genes involved in phenylpropanoid biosynthesis and extracellular processes (Figure 7C, Table 2). Finally, cluster 6 contains genes that are strongly downregulated in *med16* and upregulated in *med5ab*. These genes encode proteins involved in pollen exine formation and those that are localized to the extracellular region. Together, these data provide a basis for discovering pathways and processes that require the function of individual or multiple MED tail subunits for their regulation.

**Figure 7.**
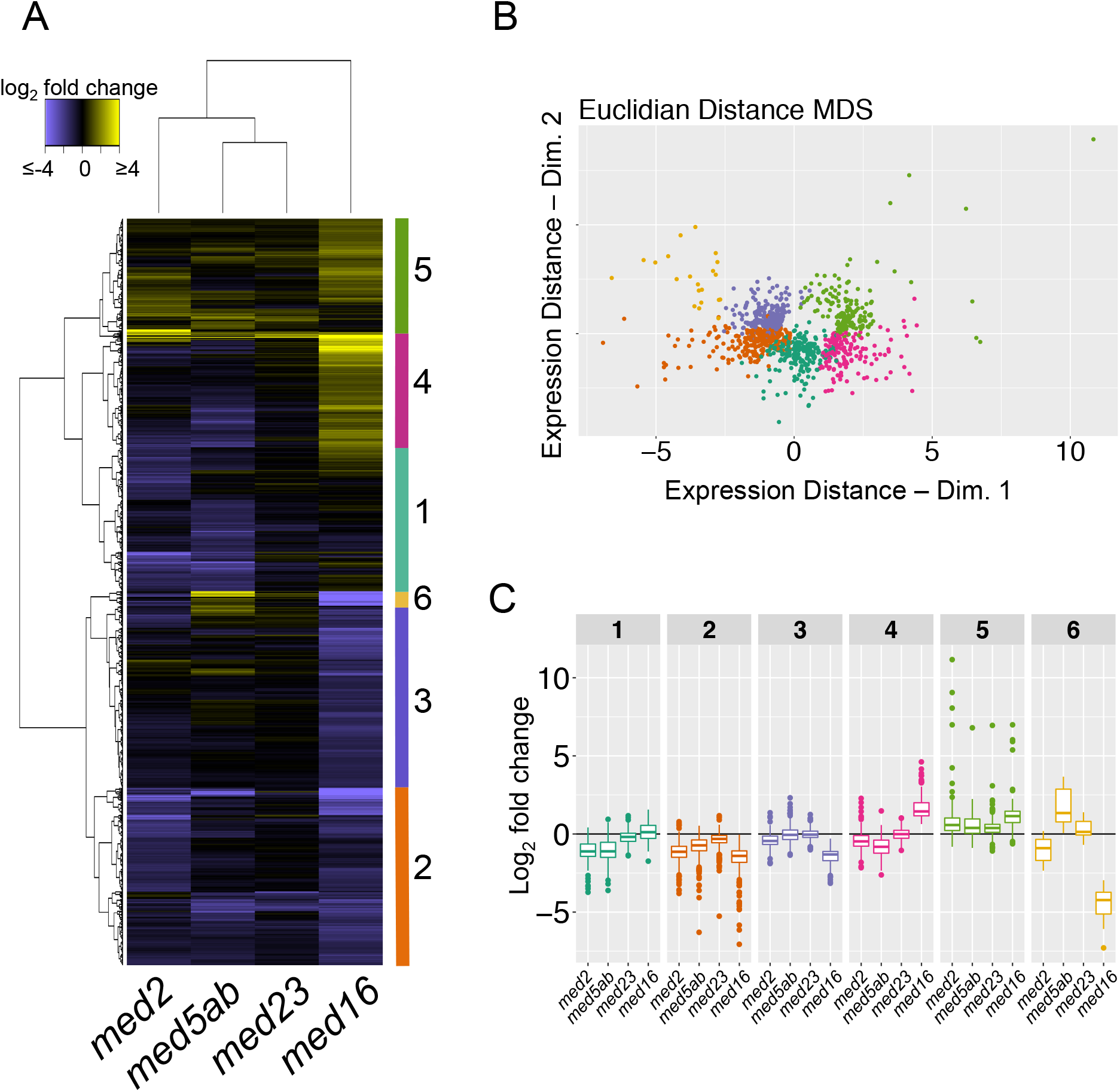
Hierarchical clustering of all genes differentially expressed in the *med* mutants. (A) Hierarchical clustering of log_2_ fold change expression values. (B) Multidimensional scaling of differentially expressed genes based on their log_2_ fold change expression values and colored by cluster membership. (C) Boxplots representing the fold-change values according to genotype and cluster membership in A and B.

**Table 2.** Gene ontology enrichment of gene clusters in Figure 7.

| Cluster total | Category | Term | Count | % | BH Pval |
| --- | --- | --- | --- | --- | --- |
| Cluster 1<br>208 | GOTERM_BP_DIRECT | GO:0050832~defense response to fungus | 16 | 7.69 | 1.50E-06 |
|  | GOTERM_BP_DIRECT | GO:0042742~defense response to bacterium | 18 | 8.65 | 2.99E-06 |
|  | GOTERM_CC_DIRECT | GO:0016021~integral component of membrane | 72 | 34.62 | 6.94E-05 |
|  | GOTERM_BP_DIRECT | GO:0010200~response to chitin | 11 | 5.29 | 8.34E-05 |
|  | GOTERM_CC_DIRECT | GO:0005886~plasma membrane | 57 | 27.40 | 1.75E-04 |
|  | GOTERM_CC_DIRECT | GO:0005576~extracellular region | 32 | 15.38 | 2.51E-04 |
|  | GOTERM_BP_DIRECT | GO:0010112~regulation of systemic acquired resistance | 5 | 2.40 | 7.60E-04 |
|  | GOTERM_BP_DIRECT | GO:0009611~response to wounding | 12 | 5.77 | 9.69E-04 |
|  | KEGG_PATHWAY | ath04626:Plant-pathogen interaction | 9 | 4.33 | 1.13E-03 |
|  | GOTERM_BP_DIRECT | GO:0009751~response to salicylic acid | 10 | 4.81 | 2.43E-03 |
|  | GOTERM_BP_DIRECT | GO:0006952~defense response | 18 | 8.65 | 2.50E-03 |
|  | GOTERM_BP_DIRECT | GO:0009753~response to jasmonic acid | 10 | 4.81 | 2.55E-03 |
|  | GOTERM_BP_DIRECT | GO:0009617~response to bacterium | 8 | 3.85 | 3.21E-03 |
|  | GOTERM_MF_DIRECT | GO:0030246~carbohydrate binding | 11 | 5.29 | 4.38E-03 |
|  | GOTERM_BP_DIRECT | GO:0012501~programmed cell death | 4 | 1.92 | 2.70E-02 |
| Cluster 2<br>257 | GOTERM_BP_DIRECT | GO:0009414~response to water deprivation | 14 | 5.45 | 1.72E-02 |
|  | KEGG_PATHWAY | ath04075:Plant hormone signal transduction | 9 | 3.50 | 4.09E-02 |
| Cluster 3<br>264 | GOTERM_CC_DIRECT | GO:0005576~extracellular region | 46 | 17.42 | 2.29E-08 |
|  | GOTERM_MF_DIRECT | GO:0046983~protein dimerization activity | 17 | 6.44 | 3.72E-07 |
|  | GOTERM_MF_DIRECT | GO:0000977~RNA polymerase II regulatory region sequence-specific DNA binding | 11 | 4.17 | 1.65E-05 |
|  | GOTERM_BP_DIRECT | GO:0000165~MAPK cascade | 8 | 3.03 | 1.10E-04 |
|  | GOTERM_BP_DIRECT | GO:0045944~positive regulation of transcription from RNA polymerase II promoter | 8 | 3.03 | 1.11E-03 |
|  | GOTERM_MF_DIRECT | GO:0033946~xyloglucan-specific endo-beta-1,4-glucanase activity | 4 | 1.52 | 1.69E-03 |
|  | GOTERM_MF_DIRECT | GO:0008794~arsenate reductase (glutaredoxin) activity | 5 | 1.89 | 1.53E-03 |
|  | KEGG_PATHWAY | ath01110:Biosynthesis of secondary metabolites | 23 | 8.71 | 1.25E-02 |
|  | GOTERM_MF_DIRECT | GO:0051537~2 iron, 2 sulfur cluster binding | 7 | 2.65 | 3.16E-03 |
|  | GOTERM_MF_DIRECT | GO:0015035~protein disulfide oxidoreductase activity | 8 | 3.03 | 3.22E-03 |
|  | KEGG_PATHWAY | ath00073:Cutin, suberine and wax biosynthesis | 4 | 1.52 | 2.23E-02 |
|  | GOTERM_MF_DIRECT | GO:0003700~transcription factor activity, sequence-specific DNA binding | 30 | 11.36 | 1.21E-02 |
| Cluster 4<br>159 | GOTERM_BP_DIRECT | GO:0009753~response to jasmonic acid | 13 | 8.18 | 9.35E-07 |
|  | GOTERM_BP_DIRECT | GO:0042742~defense response to bacterium | 16 | 10.06 | 1.31E-06 |
|  | GOTERM_BP_DIRECT | GO:0009617~response to bacterium | 11 | 6.92 | 1.62E-06 |
|  | GOTERM_BP_DIRECT | GO:0006952~defense response | 20 | 12.58 | 3.13E-06 |
|  | GOTERM_BP_DIRECT | GO:0009751~response to salicylic acid | 9 | 5.66 | 1.76E-03 |
|  | GOTERM_BP_DIRECT | GO:0009863~salicylic acid mediated signaling pathway | 5 | 3.14 | 1.90E-03 |
|  | GOTERM_BP_DIRECT | GO:0007165~signal transduction | 12 | 7.55 | 7.09E-03 |
|  | GOTERM_BP_DIRECT | GO:0050832~defense response to fungus | 9 | 5.66 | 7.76E-03 |
|  | GOTERM_BP_DIRECT | GO:0009627~systemic acquired resistance | 5 | 3.14 | 1.04E-02 |
|  | KEGG_PATHWAY | ath01110:Biosynthesis of secondary metabolites | 17 | 10.69 | 1.10E-02 |
|  | GOTERM_BP_DIRECT | GO:0055114~oxidation-reduction process | 21 | 13.21 | 1.13E-02 |
|  | GOTERM_BP_DIRECT | GO:0009695~jasmonic acid biosynthetic process | 4 | 2.52 | 2.24E-02 |
|  | GOTERM_BP_DIRECT | GO:0080027~response to herbivore | 3 | 1.89 | 2.69E-02 |
|  | GOTERM_BP_DIRECT | GO:0009611~response to wounding | 8 | 5.03 | 2.78E-02 |
|  | KEGG_PATHWAY | ath00592:alpha-Linolenic acid metabolism | 4 | 2.52 | 2.92E-02 |
|  | GOTERM_BP_DIRECT | GO:0002229~defense response to oomycetes | 4 | 2.52 | 3.13E-02 |
|  | GOTERM_CC_DIRECT | GO:0005576~extracellular region | 23 | 14.47 | 3.74E-02 |
|  | GOTERM_BP_DIRECT | GO:0009620~response to fungus | 5 | 3.14 | 4.44E-02 |
| Cluster 5<br>172 | GOTERM_CC_DIRECT | GO:0005576~extracellular region | 27 | 15.70 | 4.47E-04 |
|  | KEGG_PATHWAY | ath00940:Phenylpropanoid biosynthesis | 7 | 4.07 | 1.97E-03 |
| Cluster 6<br>21 | GOTERM_CC_DIRECT | GO:0005576~extracellular region | 11 | 52.38 | 9.89E-05 |
|  | GOTERM_BP_DIRECT | GO:0010584~pollen exine formation | 3 | 14.29 | 9.12E-03 |

### Identification of enriched transcription factor binding sites in the promoters of ***med-*** downregulated genes

DNA-bound transcription factors interact with MED subunits to regulate the expression of downstream genes. Thus, some of the changes in gene expression that are observed in the *med* mutants may stem from loss of MED-transcription factor interactions. To identify classes of transcription factors that might interact with MED2, MED5, MED16, or MED23, we assessed the enrichment of putative transcription factor binding sites in the promoters of genes that were downregulated in each of the mutants. We chose to use the downregulated genes because of the well-understood function of Mediator in gene activation. Using the Athena analysis suite we identified transcription factor binding motifs that were enriched among the promoters of *med2, med5ab*, and *med16* downregulated genes (Table 3) (O’Connor *et al*. 2005). Consistent with previously published results, both the Evening Element and the DRE/CRT motif were enriched in the promoters of *med16* downregulated genes (Knight 1999; Knight *et al*. 2008; Hemsley *et al*. 2014). There were also many other motifs enriched in these genes’ promoters. This was not surprising given the large number of downregulated genes in *med16*. Enrichment of MYC2, MYCATERD1, and MYB1AT sites was consistent with the observation that *med16* mutants show a reduced transcriptional response to ABA (Figure 6, Figure 7 Cluster 2) (Abe *et al*. 1997, 2003; Knight 1999; Simpson *et al*. 2003). There was no motif enrichment among the promoters of the *med23* downregulated genes. The W-box motif, which is bound by WRKY transcription factors, was enriched in the promoters of both *med2* and *med5ab* downregulated genes, consistent with the reduced expression of many defense-related pathway in those lines (Figure 6, Figure 7 Cluster 3) (Pandey and Somssich 2009). The coherence of these results with those of others, and with known MED functions, supports the idea that these analyses might serve as the basis for identifying novel MED-transcription factor interactions.

**Table 3.**
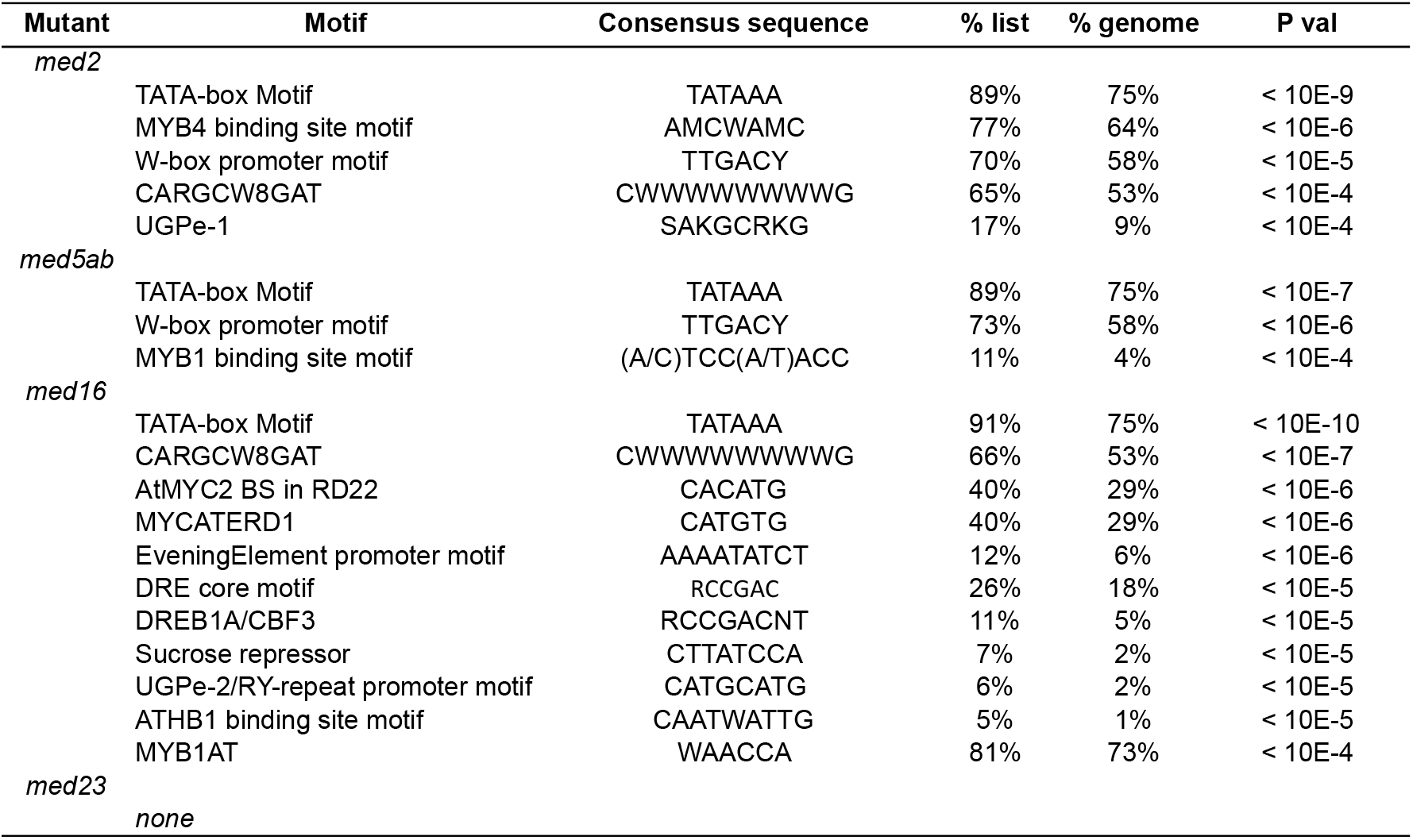
Enrichment of transcription factor binding sites in the promoters of downregulated genes

### The ***med2*** and ***med5ab*** mutants are early flowering

As previously mentioned, the *med16* mutant is late flowering (Knight *et al*. 2008) and we initially observed that the *med2* and *med5ab* mutants appeared to flower early. When we quantified this phenomenon, we found that *med2* plants flowered an average of 2.1 days earlier and with 2.6 fewer rosette leaves than wild type plants (Figure 8A and B). Similarly, *med5ab* flowered an average of 1.5 days earlier than wild type and with 2.1 fewer rosettes leaves. Consistent with the previously published results, *med16* flowered an average of 8.9 days later and with 7.3 more leaves than wild type. Additionally, *med23* plants had an average of 1.3 more leaves at the time of flowering.

**Figure 8.**
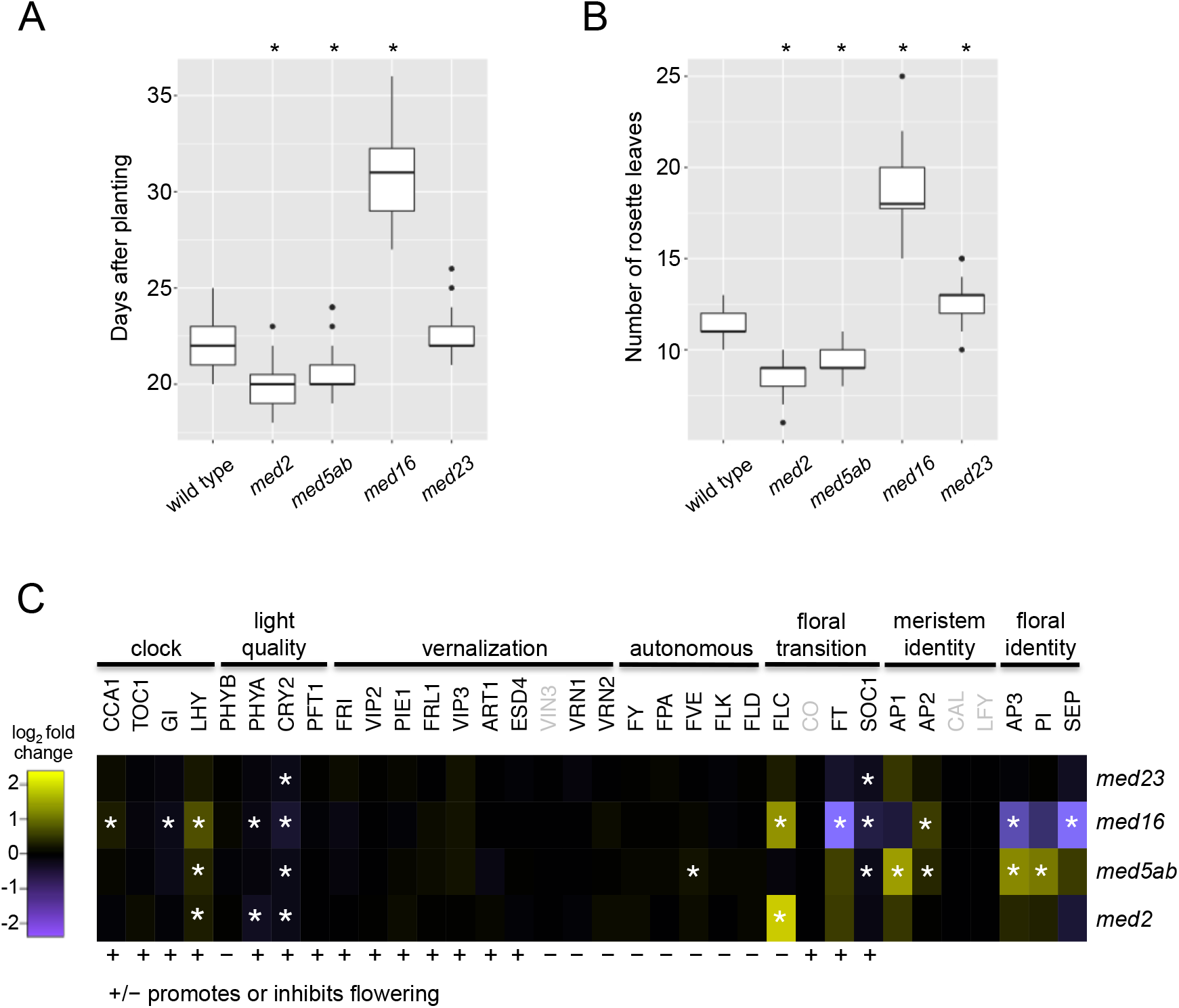
*med2* and *med5ab* are early flowering and have altered expression of flowering-related genes. (A) Days after planting and number of leaves (B) at the time that the first inflorescence reached 1 cm. Asterisks indicate *P<* 0.01 when compared to wild type (t-test, n=32-35). Boxes indicate the first quartile, the median, and the third quartile. The whiskers indicate the largest and smallest value no more than 1.5 times the interquartile range. Outliers are individually marked. (C) log_2_ fold change in expression compared to wild-type of flowering-related genes. Asterisks indicate genes with an FDR < 0.01. Genes that were not expressed are indicated in grey.

In the *med16* mutant, the late flowering phenotype was attributed to reduced expression of clock components, leading to a reduced expression of flowering genes, namely *CO* and *FT* (Knight *et al*. 2008). Although *CO* transcripts were not detectable in the samples we analyzed, expression of *FT* was strongly reduced in *med16* (Figure 8C). In addition, expression of *FLC*, a negative regulator of the floral transition (Michaels and Amasino 1999), was increased in *med16*. Examination of the major genes involved in flowering did not reveal an obvious cause for the early flowering of *med2* and *med5ab* (Figure 8C). In the case of *med2, FLC* is substantially upregulated without concomitant downregulation of it targets *SOC1* and *FT*, suggesting that *FLC* might partially require MED2 for its function in repressing the floral transition. It is also possible that the effect of *med2* and *med5ab* on flowering time is too subtle to be detected at the transcriptional level. In addition, the expression of many flowering and clock genes cycles diurnally, therefore differences in expression might be less apparent at the time we sampled the plants than at other times during the day.

### MED23 and MED5a may have tissue-specific functions

Only nine DEGs were identified in *med23*, four of which have not been characterized, lending little information as to whether MED23 has any unique functions in transcription regulation (Table 4). To explore whether MED23 might play a more predominant role in other organs or in particular tissues, we used the Arabidopsis eFP browser to compare the expression of the *MED23, MED5a, MED5b* and *MED16* (data for *MED2* was not available) during the development of different organs (Figure 9A) and in different tissues (Figure 9B) (Winter *et al*. 2007). *MED23* was expressed in all organs, but was expressed more strongly in seeds, flowers, roots, and shoots than in leaves (Figure 9A). *MED23* also showed substantial expression in mature pollen, whereas the other *MED* genes did not. Most striking, however, was the strong expression of *MED23* in the shoot apical meristem (Figure 11B, “Peripheral zone”, “Central zone”, “Rib meristem”). These data suggest that *MED23* might have specific functions in meristematic or reproductive development. This hypothesis is strengthened by the observation that the floral specification gene *AGAMOUS (AG)* is upregulated in *med23* (Table 4), and that several other genes involved in embryo, floral, or meristem development are co-expressed with *MED23* (Table 5) (ATTED-II v8.0, Aoki et al., 2016). The eFP data also showed that MED5a is more highly expressed than MED5b during most developmental stages, and has a much higher level of expression in guard cells than the other MED subunits we examined.

**Figure 9.**
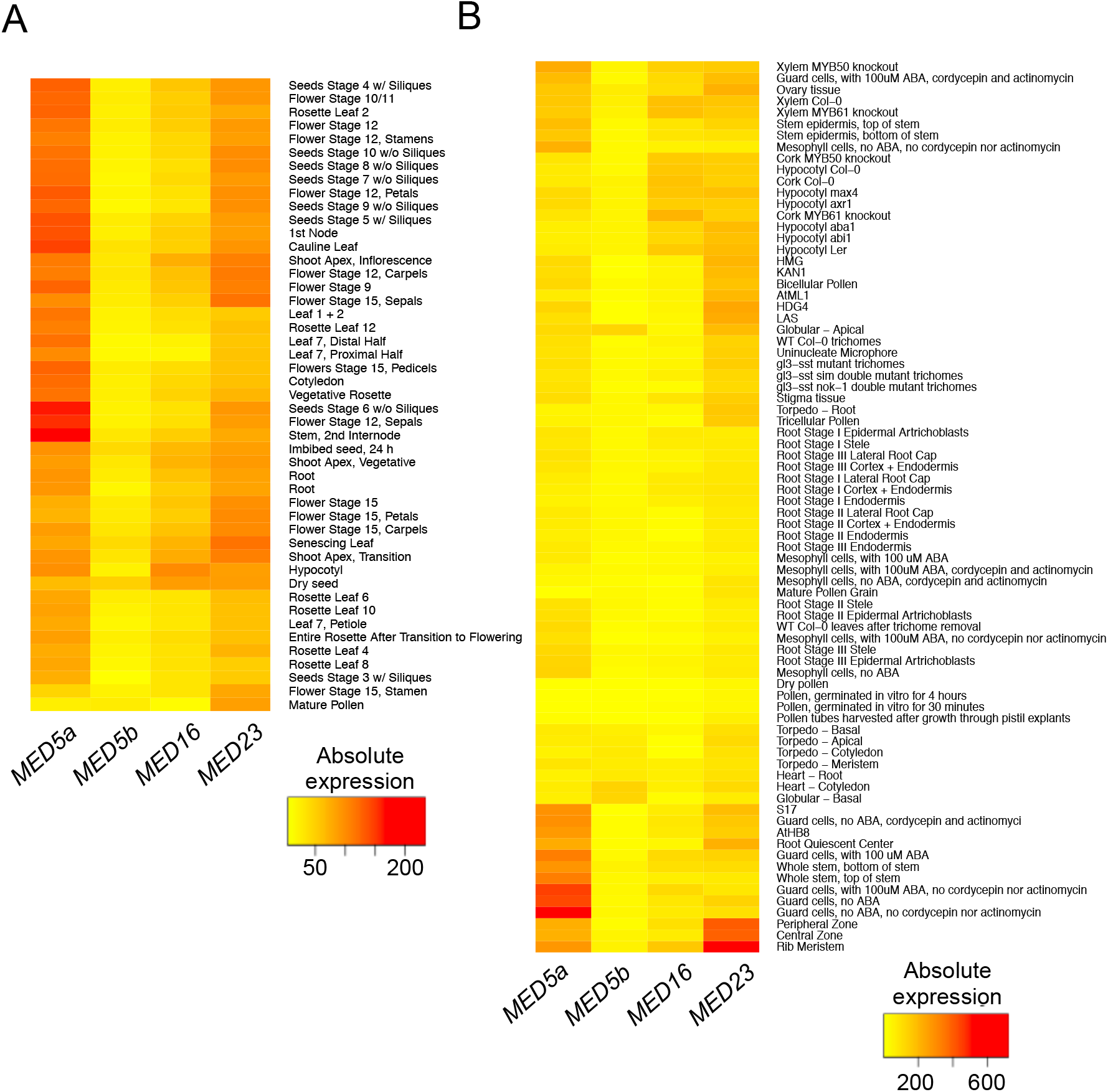
Expression of the *MED5ab, MED5b, MED16 and MED23* across development of different organs and in different tissues. The (A) “Development” and (B) “Tissue” datasets were retrieved from the Arabidopsis eFP browser.

**Table 4.** Genes that are differentially expressed in *med23* compared to wild type

| AGI | log2FC | BH P val | Gene description |
| --- | --- | --- | --- |
| AT5G35935 | 6.94 | 1.60E-240 | copia-like retrotransposon family |
| AT4G08093 | 3.09 | 3.38E-03 | expressed protein |
| AT3G30122 <sup>a</sup> | 1.60 | 2.38E-15 | expressed protein |
| AT5G28335 | 1.59 | 1.74E-03 | gypsy-like retrotransposon family |
| AT2G01008 <sup>a</sup> | 1.58 | 2.87E-07 | unknown protein, best match MEE38 |
| AT3G41979 | 1.56 | 1.82E-14 | 5S rRNA |
| AT4G07850 | 1.46 | 1.79E-03 | gypsy-like retrotransposon family |
| AT2G16367 | 1.40 | 6.46E-03 | defensin-like (DEFL) family protein |
| AT3G44765 | 1.37 | 8.90E-03 | other RNA |
| AT2G05914 | 1.37 | 5.86E-03 | Natural antisense transcript overlaps with AT2G05915 |
| AT3G44970 <sup>a</sup> | 1.24 | 3.12E-11 | Cytochrome P450 superfamily protein |
| AT1G30760 | 1.16 | 6.41E-03 | FAD-binding Berberine family protein |
| AT3G22415 <sup>a</sup> | 1.06 | 9.44E-03 | unknown protein |
| AT3G19550 <sup>a</sup> | 1.03 | 1.30E-03 | unknown protein |
| AT4G18960 <sup>a</sup> | 1.02 | 2.98E-06 | AGAMOUS (AG) |
| AT1G23230 <sup>a</sup> | -5.26 | 0.00E+00 | MED23 |
| AT1G35210 | -2.35 | 3.30E-07 | unknown protein |
| AT5G51190 | -2.15 | 2.93E-07 | Integrase-type DNA-binding superfamily protein |
| AT4G17490 | -2.07 | 1.28E-06 | ethylene responsive element binding factor 6 (ERF6) |
| AT4G24570 | -2.06 | 2.92E-11 | dicarboxylate carrier 2 (DIC2) |
| AT2G25735 | -1.82 | 4.05E-10 | unknown protein |
| AT3G44260 | -1.74 | 2.73E-13 | Polynucleotidyl transferase, ribonuclease H-like |
| AT3G29000 | -1.71 | 6.46E-03 | Calcium-binding EF-hand family protein |
| AT5G27420 | -1.67 | 1.96E-04 | carbon/nitrogen insensitive 1 (CNI1) |
| AT5G04340 | -1.64 | 1.68E-04 | zinc finger of Arabidopsis thaliana 6 (ZAT6) |
| AT5G61600 | -1.62 | 7.14E-10 | ethylene response factor 104 (ERF104) |
| AT1G07135 | -1.61 | 2.06E-04 | glycine-rich protein |
| AT1G27730 | -1.58 | 8.85E-06 | salt tolerance zinc finger (STZ) |
| AT5G47230 | -1.54 | 1.29E-08 | ethylene responsive element binding factor 5 (ERF5) |
| AT5G56320 | -1.50 | 3.77E-08 | expansin A14 (EXPA14) |
| AT1G66090 | -1.45 | 4.66E-09 | Disease resistance protein (TIR-NBS class) |
| AT1G74290 <sup>a</sup> | -1.40 | 7.78E-06 | alpha/beta-Hydrolases superfamily protein |
| AT1G53480 | -1.38 | 2.19E-12 | mto 1 responding down 1 (MRD1) |
| AT4G23810 | -1.34 | 2.41E-05 | WRKY family transcription factor (WRKY53) |
| AT3G30720 | -1.33 | 5.52E-07 | qua-quine starch (QQS) |
| AT5G45340 | -1.26 | 7.79E-05 | CYP707A3 |
| AT2G33830 | -1.23 | 1.29E-05 | Dormancy/auxin associated family protein |
| AT5G23240 | -1.20 | 1.85E-06 | DNAJ heat shock N-terminal domain-containing protein |
| AT3G51860 | -1.17 | 2.28E-03 | cation exchanger 3 (CAX3) |
| AT2G38470 | -1.16 | 4.15E-08 | WRKY DNA-binding protein 33 (WRKY33) |
| AT2G01010 | -1.15 | 3.81E-04 | 18S rRNA |
| AT5G59820 | -1.13 | 6.84E-03 | C2H2-type zinc finger family protein (RHL41) |
| AT3G55980 <sup>a</sup> | -1.11 | 7.92E-12 | salt-inducible zinc finger 1 (SZF1) |
| AT2G47260 | -1.11 | 5.75E-13 | WRKY DNA-binding protein 23 (WRKY23) |
| AT3G16720 | -1.10 | 2.43E-09 | TOXICOS EN LEVADURA 2 (ATL2) |
| AT4G29780 | -1.08 | 3.69E-04 | unkown protein |
| AT5G26920 | -1.02 | 1.96E-04 | Cam-binding protein 60-like G (CBP60G) |
| AT2G24600 | -1.00 | 7.72E-05 | Ankyrin repeat family protein |

<sup>a</sup> Genes that are differentially expressed in *med23* but not the other med mutants.

**Table 5.** Top 40 genes coexpressed with *med23* according to mutual rank

| AGI | Alias | Function | Mutual Rank<sup>c</sup> |
| --- | --- | --- | --- |
| At1g02080<sup>a</sup> | transcription | transcription regulators | 6.9 |
| At1g48090 | calcium-dependent lipid-binding | calcium-dependent lipid-binding family protein | 7.1 |
| At1g80070<sup>a</sup> | SUS2 | Pre-mRNA-processing-splicing factor | 7.9 |
| At5g58410 | HEAT/U-box | HEAT/U-box domain-containing protein | 13 |
| At4g39850 | PXA1 | peroxisomal ABC transporter 1 | 13.8 |
| At3g13330 | PA200 | proteasome activating protein 200 | 16.2 |
| At3g02260<sup>b</sup> | UMB1 | auxin transport protein (BIG) | 17.2 |
| At1g50030<sup>a,b</sup> | TOR | target of rapamycin | 17.3 |
| At5g23110 | Zinc finger | Zinc finger, C3HC4 type (RING finger) family protein | 20.4 |
| At4g01290 | chorismate synthase | chorismate synthase | 20.5 |
| At2g26780 | ARM repeat | ARM repeat superfamily protein | 22.4 |
| At3g27670 | RST1 | ARM repeat superfamily protein | 22.9 |
| At2g17930 | Phosphatidylinositol 3- and 4-kinase | Phosphatidylinositol 3- and 4-kinase family protein with FAT domain | 23.8 |
| At1g20960<sup>a</sup> | emb1507 | U5 small nuclear ribonucleoprotein helicase, putative | 25.1 |
| At1g54490<sup>a</sup> | XRN4 | exoribonuclease 4 | 26.3 |
| At2g41700 | ABCA1 | ATP-binding cassette A1 | 29.1 |
| At1g15780 | NRB4/MED15a | Mediator subunit 15a | 29.7 |
| At5g61140<sup>a</sup> | helicase | U5 small nuclear ribonucleoprotein helicasea | 30.7 |
| At5g51340 | Tetratricopeptide repeat (TPR)-like | Tetratricopeptide repeat (TPR)-like superfamily protein | 31.4 |
| At3g57570 | ARM repeat | ARM repeat superfamily protein | 31.9 |
| At5g15680 | ARM repeat | ARM repeat superfamily protein | 33 |
| At4g00450<sup>b</sup> | MED12 | RNA polymerase II transcription mediators | 33.5 |
| At3g15880<sup>b</sup> | WSIP2 | WUS-interacting protein 2 | 34.3 |
| At3g51050 | FG-GAP repeat | FG-GAP repeat-containing protein | 37.5 |
| At3g16830<sup>b</sup> | TPR2 | TOPLESS-related 2 | 40.3 |
| At3g50590 | Transducin/WD40 repeat-like | Transducin/WD40 repeat-like superfamily protein | 45.2 |
| At1g72390 | PHL | Phytochrome-dependent late-flowering | 45.5 |
| At3g60240 | EIF4G | eukaryotic translation initiation factor 4G | 48 |
| At5g16280 | Tetratricopeptide repeat (TPR)-like | Tetratricopeptide repeat (TPR)-like superfamily protein | 48.6 |
| At3g08850<sup>b</sup> | RAPTOR1B | HEAT repeat ;WD domain, G-beta repeat protein protein | 49 |
| At1g55325<sup>b</sup> | MAB2/MED13 | RNA polymerase II transcription mediators | 49.9 |
| At5g47010<sup>a</sup> | UPF1 | RNA helicase, putative | 50.2 |
| At3g33530 | Transducin | Transducin family protein / WD-40 repeat family protein | 52 |
| At2g32730 | proteasome | 26S proteasome regulatory complex, Rpn2/Psmd1 subunit | 52.5 |
| At5g65750 | 2-oxoglutarate dehydrogenase | 2-oxoglutarate dehydrogenase, E1 component | 54 |
| At3g07160 | GSL10 | glucan synthase-like 10 | 54.7 |
| At2g28290<sup>b</sup> | SYD | P-loop containing nucleoside triphosphate hydrolases superfamily protein | 55.1 |
| At2g33730<sup>a</sup> | hydrolase | P-loop containing nucleoside triphosphate hydrolases superfamily protein | 55.6 |
| At5g51660<sup>a</sup> | CPSF160 | cleavage and polyadenylation specificity factor 160 | 58.8 |
| At3g50380 | DUF1162 | Protein of unknown function (DUF1162) | 59 |

<sup>a</sup> Genes annotated as being involved in RNA processing
<sup>b</sup> Genes annotated as being involved in embryo, floral or meristem development
<sup>c</sup> Based on ATTED-II data set Ath-m.v15-08

### MED tail mutants might affect alternative mRNA processing

Many genes involved in RNA processing are co-expressed with *MED23* (Table 5). In humans, MED23 interacts with mRNA processing factors and is required for the alternative splicing and polyadenylation of a significant number of transcripts (Huang *et al*. 2012). To determine if MED23 or the other MED subunits examined here might be involved in alternative splicing in Arabidopsis, we queried our RNAseq data for differential splicing events using the diffSpliceDGE function in edgeR (Robinson *et al*. 2010). To detect alternative exon usage, diffSpliceDGE compares the log fold change of individual exons to that of the gene as a whole. Using an FDR cutoff of 0.05, we detected a handful of alternatively spliced (AS) transcripts in each of the mutants, with the most being found in *med23* (Figure 10A). The vast majority of these were not differentially expressed at the level of the whole gene. GO-term enrichment analysis of the AS transcripts found in each mutant showed that genes encoding ribosomal proteins, membrane proteins, chloroplast localized proteins, vacuolar proteins, and cell wall proteins were enriched in all four mutants (FDR < 0.05). In addition, the UniProt keyword “alternative splicing” was also enriched in all four lists, indicating that many of these transcripts have previously been shown to be alternatively spliced. Of the approximately 30 alternative splicing events that we examined, all but one occurred at the 5’ or 3’ end of the gene (e.g. Figure 10B), with many occurring within the untranslated region. They also all exhibited relatively small fold-changes, such that they could not be identified from coverage maps by eye. Together, these results suggest that these MED subunits, particularly MED23, might influence alternative RNA processing, either directly or indirectly.

**Figure 10.**
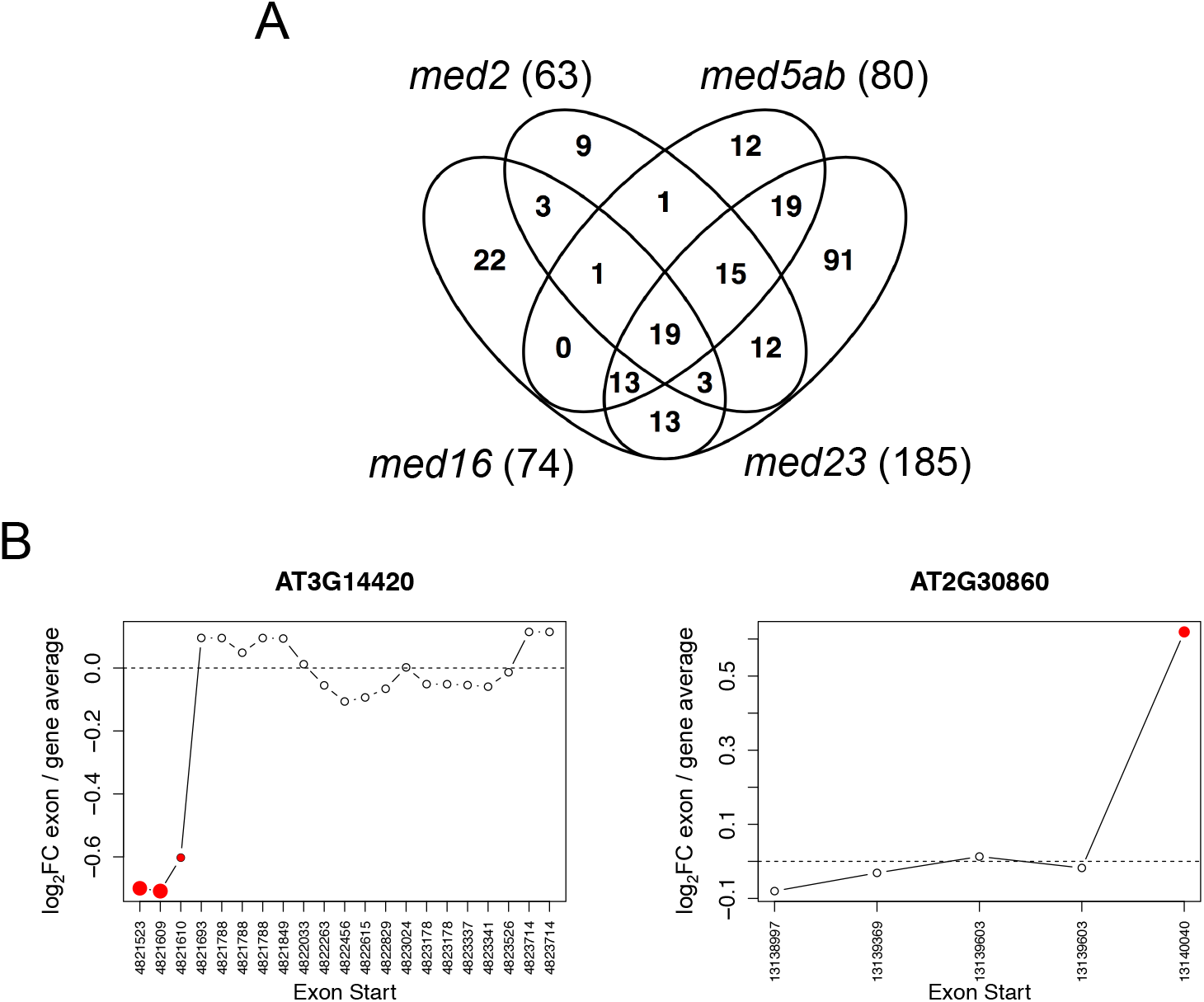
Alternative splicing occurs in the 5’ and 3’ ends of genes in the *med* mutants. (A) Number of alternatively spliced transcripts in the *med* mutants (FDR < 0.05). (B) Two examples of transcripts that are alternatively spliced in *med23*. Log_2_ fold change in the expression of individual exons compared to that of the entire gene. Significant exons are highlighted in red.

One of the “alternative splicing” events appeared very different from the rest. *AT1G64790* was detected as an alternatively spliced transcript in *med2* and *med23* because of a large number of reads that mapped to a region spanning the first and second introns of the gene that were not present in wild type (Figure 11). According to the Araport11 annotation of the Arabidopsis genome, this region produces a cluster of 24 nt small RNAs. It has also been shown to be differentially methylated in the C24 and Ler ecotypes, and undergoes

**Figure 11.**
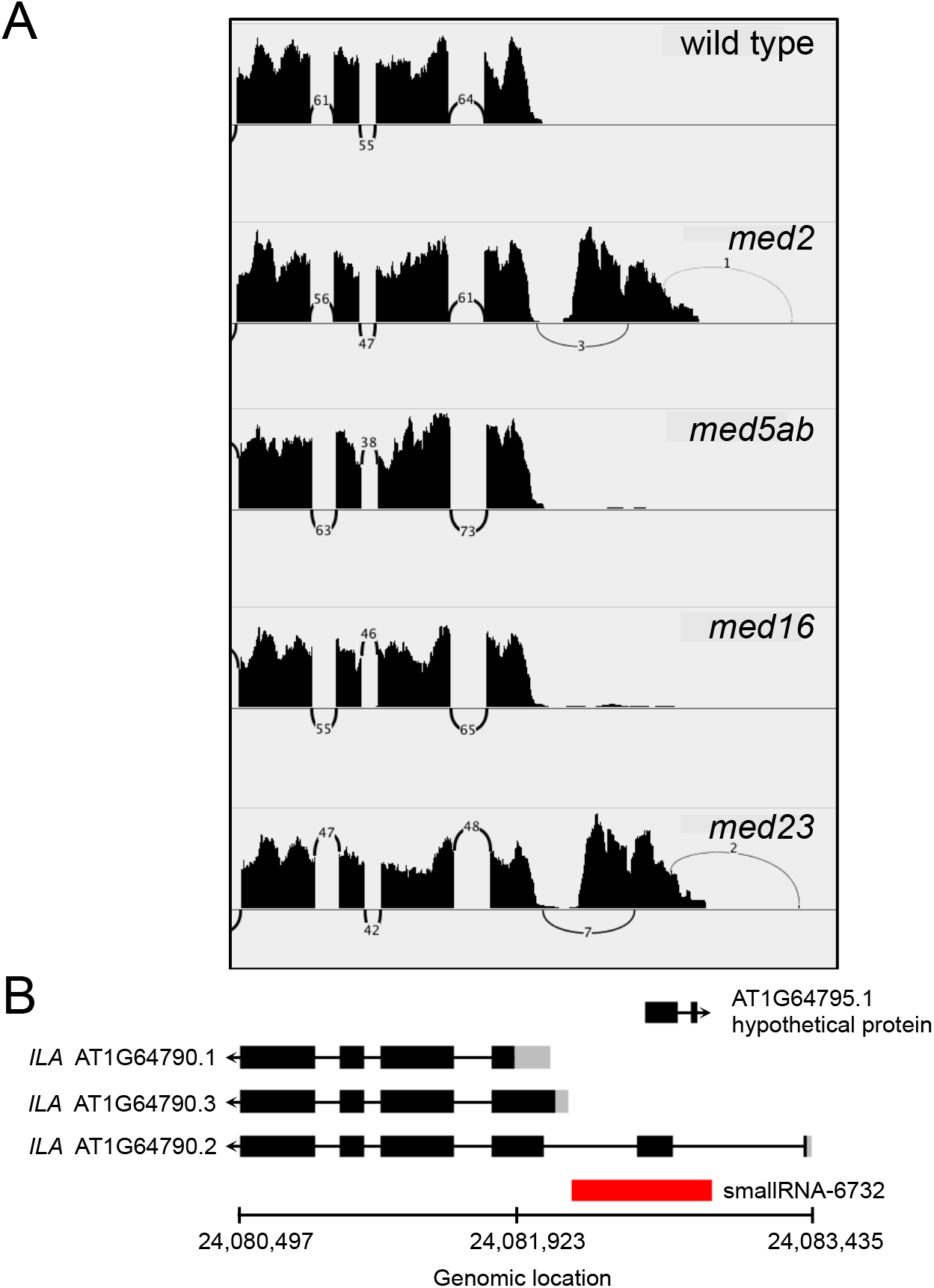
A region that undergoes transcriptional gene silencing is derepressed in *med2* and *med5ab*. (A) Read coverage and number of intron-spanning reads across (B) a region of chromosome 1, which includes a portion of the *ILYTHIA* gene, a hypothetical protein and a smallRNA. Exons are indicated as black rectangles, UTRs are in grey. Coverage in (A) is from individual wild-type or mutant samples. transchromosomal methylation in F2 hybrids (Greaves *et al*. 2014). Mediator has previously been shown to be required for RNA-directed DNA methylation of repeats and transposons (Kim and Chen 2011). The derepression of this region in the *med2* and *med23* mutants suggests that they are specifically required for this process at some loci.

## DISCUSSION

Arabidopsis is largely robust to perturbation of many Mediator complex subunits. Unlike mice, in which all MED knockouts tested have proved to be embryonic lethal, most of the Arabidopsis *MED* mutants studied to date grow well enough in controlled environments that they are fertile (Yin and Wang 2014; Buendía-Monreal and Gillmor 2016). This makes Arabidopsis uniquely suited to studying the effects of disruption of the complex in a developing, multicellular eukaryote. Many studies of Arabidopsis MED mutants have examined the effects of disruption of one or a few MED subunits on a limited number pathways or genes. In the present study, we sought to gain a broader understanding of the function of the Arabidopsis tail module and the relative contributions of its subunits to genome-wide transcription by comparing the effects of mutations in four different MED tail subunits—MED2, MED5a/b, MED16, and MED23—on the transcriptome.

The T-DNA mutants studied here all develop normally, enabling our analysis of changes in gene expression to be unencumbered by changes that might arise due to altered morphology (Figures 2 and 8). We did, however, observe that *med16* flowered late, in accordance with previous reports (Knight et al. 2008), and that *med2* and *med5* flowered early (Figures 8A and 8B). In addition to *med16*, mutations in eight other MED subunits cause Arabidopsis to flower late (Reviewed in Yang et al. 2015). The *med2* and *med5ab* mutants are unique in that they are the only MED mutants reported to date that cause plants to flower early. Given the large network of genes that impinge on flowering time, and the broad involvement of Mediator in transcriptional regulation, it is not surprising that so many MED mutants affect flowering time. Although our gene expression analysis pointed to a potential reason for the early flowering of *med2*, additional studies will be required to determine the mechanistic cause. At the time that rosettes were sampled for RNAseq analysis, some plants had formed an apical bud. This may explain why genes related related to pollen exine formation appeared to be upregulated in *med5ab* and downregulated in med16 (Figure 7A, Table 2, Cluster 6)

In the collection of MED mutants we examined, relatively few genes passed our criteria for differential expression (Figure 4). We found that, although the mutants shared many differentially expressed genes, there were also a large number of genes that were uniquely differentially expressed in each mutant. Genes that were upregulated in all four mutants showed an enrichment of genes encoding extracellular proteins, as well as phenylpropanoid related genes (Figure 7A, Cluster 5), consistent with their ability to rescue the phenylpropanoid-deficient mutant *ref4-3* (Dolan et al. 2017). Many of the genes that were altered in the mutants were related to abiotic or biotic stress, in which Mediator is known to play a major role (Samanta and Thakur 2015b). The *med16* and *med5ab* mutants were the most different from one another, showing opposite regulation of many of the same genes (Figures 5B and 7A). Conversely, we observed a strong correlation between the gene expression profiles of *med5ab* and *med23* (Figure 5B). This finding is consistent with our previous observation that both *med5ab* and *med23* have higher levels of sinapoylmalate (Dolan et al. 2017) and suggests a broad functional link between the two subunits, possibly mediated by a close physical association within the complex. This close association is also supported by the observation that knocking out *med23* in the *MED5b* mutant *ref4-3* strongly and specifically suppresses the transcriptional and phenotypic effects of *ref4-3* (Dolan et al. 2017). Our data also suggest that MED2 plays a more general role in gene regulation than some of the other MED subunits, as only a small number of pathways were significantly enriched in the *med2* mutant, despite the substantial number of genes that are differentially expressed in that line (Figures 4 and 6).

As we previously reported, the *med16* mutant is different from the other *med* mutants investigated here, in that a large number of genes are upregulated in the mutant, consistent with what has been observed in the yeast (Chen *et al*. 1993; Covitz *et al*. 1994; Jiang and Stillman 1995). What was more surprising was that the genes that were upregulated in *med16* were associated with defense pathways, including those controlled by salicylic acid and jasmonic acid (Figure 7, Table 2, Cluster 4). MED16 has been extensively reported as being a positive regulator of both SA and JA-mediated defense (Wathugala *et al*. 2012; Zhang *et al*. 2012, 2013, Wang *et al*. 2015, 2016). Given the existence of numerous positive and negative regulators of these pathways, close inspection of the identity and function of these genes will be required to determine how these findings fit with known role of MED16 in defense response pathways. Additionally, many of these genes are downregulated in *med5ab* (Figure 7, Cluster 4), suggesting a possible antagonistic or epistatic relationship between MED5a/b and MED16 in the expression of defense response genes.

MED23 is one of several subunits that are conserved in metazoans and plants, but not in Saccharomyces (Bourbon 2008). In humans, MED23 plays a variety of important roles, including promoting transcription elongation, alternative splicing, and ubiquitination of histone H2B (Huang *et al*. 2012; Wang *et al*. 2013; Yao *et al*. 2015). Aside from our previous report, the role of MED23 in transcription regulation in plants has yet to be investigated. Our data suggest that MED23 does not play a major role in Arabidopsis rosettes under normal growth conditions. Examination of the expression of *MED23* in different organs and tissues, as well as the genes that are co-expressed with *MED23*, suggested that MED23 might function in reproductive or meristem development. Two of the genes co-expressed with *MED23, MED12 and MED13*, encode subunits of the Mediator kinase module. MED12 and MED13 play a transient role in early embryo patterning and development, and similar to MED23, are expressed most strongly in the shoot apical meristem (Gillmor *et al*. 2010). Together, these observations suggest that MED23 might function together with MED12 and MED13 in embryo development, particularly in establishing the shoot apical meristem.

We also discovered evidence of a conserved role for MED23 in alternative splicing, in that a number of RNA processing factors are co-expressed with *MED23* and that more alternative transcripts were produced in *med23* than in the other mutants (Table 4, Figure 10A). All of the alternative splicing events that we examined occurred at either 5’ or 3’ ends of the genes. GO-term analysis of these genes showed an enrichment of genes encoding membrane proteins or proteins localized to different cellular compartments. Alternative splicing of N– or C-terminal exons can affect where proteins are targeted by changing the inclusion of signal peptides or transmembrane helices (Davis *et al*. 2006; Dixon *et al*. 2009; Lamberto *et al*. 2010; Kriechbaumer *et al*. 2012; Remy *et al*. 2013). In addition, alternative UTRs can affect transcript stability and translation efficiency (Reddy *et al*. 2013). Biochemical validation will be required to determine whether these transcripts truly undergo alternative splicing in the *MED* mutants, and if so, what consequences they have on protein function or localization. Two major mechanisms have been proposed by which Mediator might affect splicing. In the “recruitment model”, Mediator and Pol II impact splicing by directly interacting with splicing factors to facilitate their recruitment to the transcription machinery (Merkhofer *et al*. 2014). MED23 has been shown to function in this way in HeLa cells by interacting with and promoting the recruitment of the splicing factor hnRNPL (Huang *et al*. 2012). Alternatively, Mediator might affect splicing by altering the rate of the transcription elongation, the so-called “kinetic model” of co-transcriptional splicing (Donner *et al*. 2010; Takahashi *et al*. 2011; Wang *et al*. 2013).

In the course of our alternative splicing analysis, we discovered that in *med2* and *med23* a region that appears to undergo transcriptional gene silencing (TGS) was derepressed (Figure 11, Greaves *et al*. 2014). Previously, mutation of Arabidopsis MED17, MED18 or MED20a was shown to disrupt TGS at type II loci by reducing the efficiency with which Pol II is recruited, causing reduced production of the long noncoding scaffold RNAs required for the recruitment of Pol V (Kim and Chen 2011). These MED subunits were also shown to be required for TGS at some type I loci, which are not known to require Pol II for silencing, but the mechanism by which they are required is unknown. Whether MED2 and MED23 function similarly remains to be seen.

This study is the first to present a side-by-side comparison of the effects of multiple Arabidopsis *med* mutants on global gene expression. Importantly, these data begin to unravel the complex network of interactions within Mediator that are required for the regulation of different genes and pathways and they suggest a number of potential avenues for future investigation.

## METHODS

### Plant Materials and Growth

*Arabidopsis thaliana* (ecotype Columbia-0) was grown in Redi-earth Plug and Seedling Mix (Sun Gro Horticulture, Agawam, MA) at a temperature of 23°C, under a long-day (16 hr light/8 hr dark) photoperiod with a light intensity of 100 μE m^-2^ s^-1^. Seeds were planted nine per 4” x 4” pot and held for two days at 4°C before transferring to the growth chamber.

Salk insertion lines were obtained from the Arabidopsis Biological Resource Center (Ohio State University) unless otherwise noted. The insertion lines used in this study include: *med5b-1 /ref4-6* (SALK_037472), *med5a-1/rfr1-3* (SALK_011621) (Bonawitz *et al*. 2012), *med2-1* (SALK_023845C) (Hemsley *et al*. 2014), *sfr6-2* (SALK_048091) (Knight *et al*. 2009). The *med2-1*, and *med23-4* mutants were provided to us by Dr. Tesfaye Mengiste (Department of Botany and Plant Pathology, Purdue University). The *med16-1/sfr6-2* mutant was provided by Dr. Zhonglin Mou (Department of Microbiology and Cell Science, University of Florida). Salk lines were genotyped as previously described (Dolan et al. 2017).

### Determination of Flowering Time

Pots were randomized within the growth chamber to minimize positional effects on growth. The number of rosette leaves was counted on the first day that the inflorescence reached or exceeded 1 cm and that day was recorded as the day of flowering.

### RNA Extraction and Whole Transcriptome Sequencing

Samples were collected for whole transcriptome sequencing (RNAseq) 20 days after planting, 6.5 hours after subjective dawn. For each biological replicate, five whole rosettes were pooled from five different pots with randomized locations and immediately flash frozen in liquid nitrogen. Samples were then stored at −80°C until RNA extraction. For RNA extraction, the pooled rosettes were ground to a powder under liquid nitrogen using a chilled mortar and pestle. Approximately 80 mg of ground tissue was then transferred to an Eppendorf tube for RNA extraction using the RNEasy Plant Mini kit from Qiagen (Qiagen, Chatsworth, CA). Total RNA was submitted to the Purdue Genomics Core Facility (Purdue University) for purification of polyA+ RNA, library construction, and sequencing. All samples were dual-barcoded, pooled, and loaded onto 4 sequencing lanes. Paired-end, 100 bp sequencing was performed by an Illumina HiSeq2500 machine run in “rapid” mode (Illumina, San Diego, CA). Read mapping was performed by the Purdue Genomics Core using the TAIR10 genome build and Tophat v. 2.1.0 (Trapnell *et al*. 2009). Transcriptome data has been deposited with the Gene Expression Omnibus under accession GSE95574.

### Statistical Analysis of RNAseq Data

RNAseq data was acquired as described previously (Dolan et al. 2017). Briefly, digital gene expression (counts) for every exon was determined using the HTSeq-count program with the intersection “nonempty” option (Anders *et al*. 2015). Counts were summarized by gene ID. The edgeR program was used for differential gene expression analysis (Robinson *et al*. 2010). The analysis began with a count table comprising 33,602 genes. Genes expressed at low levels were filtered out by removing any genes for which there was not at least 1 count per million in at least four of the samples. This resulted in a list of 18,842 expressed genes. The exact test for the negative binomial distribution was then used to identify genes that were differentially expressed in the *med* mutants compared to wild type (FDR < 0.01) (Robinson and Smyth 2008). The results of these analyses are available in Supplemental File S1. Gene ontology enrichment was performed using DAVID v6.8 (Huang *et al*. 2008). All genes that were expressed in our data set were used as the set of background genes for enrichment testing. GO terms were considered enriched if the associated Benjamini-Hochberg adjusted *P* value was less than 0.05. Where noted, redundant GO terms were removed for the purposes of reporting.

Alternative splicing analysis was performed using the procedure provided in the edgeR package (Robinson *et al*. 2010). Testing was performed between each *med* mutant and wild type using the diffSpliceDGE function (Lun *et al*. 2016). Simes’ method was used to convert exon-level *P* values to genewise *P* values. Genes with an FDR < 0.05 were considered as having alternatively spliced transcripts.

The Athena analysis suite was used to identify and test for enrichment of transcription factor binding motifs within 1000 bp of the transcription start site of each gene. Motifs were considered enriched if the associated *P* value was less than 10E-4 (cutoff recommended by the Athena developers based on a Bonferroni correction).

## Data and Reagent Availability

Gene expression data has been deposited with the Gene Expression Omnibus under accession GSE95574. Supplemental File 1 has been uploaded to the GSA Figshare portal. File S1 contains the results of our differential expression analysis. Arabidopsis MED T-DNA lines are available upon request.

## ACKNOWLEDGEMENTS

This work was supported by the U.S. Department of Energy, Office of Science, Office of Basic Energy Sciences, Chemical Sciences, Geosciences, and Biosciences Division under Award DE-FG02-07ER15905 and Stanford University’s Global Climate and Energy Project (GCEP).

